# Cell-based assay to determine Type 3 Secretion System translocon assembly in *Pseudomonas aeruginosa* using split luciferase

**DOI:** 10.1101/2023.06.22.546099

**Authors:** Hanling Guo, Emily Geddes, Timothy J. Opperman, Alejandro P. Heuck

**Author notes:** Corresponding Authors (A.P.H.); (T.J.O.).

## Abstract

Multidrug resistant *Pseudomonas aeruginosa* poses a serious threat to hospitalized patients. This organism expresses an arsenal of virulence factors that enables it to readily establish infections and to disseminate in the host. The Type III secretion System (T3SS) and its associated effectors play a crucial role in the pathogenesis of *P. aeruginosa,* making them attractive targets for the development of novel therapeutic agents. The T3SS translocon, comprised of PopD and PopB, is an essential component of the T3SS secretion apparatus. In the properly assembled translocon, the N-terminus of PopD protrudes into the cytoplasm of the target mammalian cell, which can be exploited as a molecular indicator of functional translocon assembly. In this manuscript, we describe a novel whole-cell-based assay that employs the split NanoLuc luciferase detection system to provide a readout for translocon assembly.

The assay demonstrates a favorable signal/noise ratio (17.9) and robustness (z’=0.73), making it highly suitable for high-throughput screening of small molecule inhibitors targeting T3SS translocon assembly.

---

*Pseudomonas aeruginosa* is an opportunistic Gram-negative pathogen that is known to cause severe, healthcare-associated infections, including hospital-acquired and ventilator-associated bacterial pneumonia [1, 2]. Despite appropriate antibiotic treatment, mortality rates can reach up to ∼70% for patients with ventilator-associated bacterial pneumonia [3]. Antibacterial therapy options for *P. aeruginosa* infections are limited due to the intrinsic resistance of the species, resulting from an impermeable outer membrane and active efflux pumps [4, 5]. Further, with the rise of acquired antibiotic resistance, some strains of *P. aeruginosa* that are resistant to almost all classes of antibiotics have emerged [6, 7]. As such, innovative strategies for the treatment of *P. aeruginosa* infections are greatly needed. One non-traditional approach has focused on targeting virulence mechanisms, as inhibition of virulence has been shown to reduce morbidity and improve the host immune response to infection [8-10]. In *P. aeruginosa*, the Type 3 Secretion System (T3SS) is one of the most clinically important virulence mechanisms [11]. During acute respiratory infections, strains that express a functional T3SS intoxicate neutrophils that infiltrate the site of infection, resulting in localized immunosuppression during pneumonia [12], which enables *P. aeruginosa* to effectively establish infection in the lung.. Importantly, higher rates of recurrent infections, morbidity, and mortality are associated with clinical isolates that express the T3SS [13]. The development of T3SS inhibitors will provide a significant therapeutic benefit for these hard-to-treat infections by rescuing innate immune function [14-16].

The T3SS facilitates the secretion and translocation of bacterial effector proteins from the cytosol of the bacterium into the cytosol of the host cell [17-19]. In *P. aeruginosa* more than 42 coordinately regulated genes encode the secretion apparatus, the effector proteins (ExoS, ExoU, ExoT, and ExoY), a translocon, and various regulatory factors [reviewed in 19, see Figure 1a]. The secretion apparatus is comprised of a cytosolic platform, an export apparatus, a multiring basal body that spans the inner membrane, peptidoglycan layer, and outer membrane of bacteria, a hollow needle that is anchored in the membrane through the basal body and extends into the extracellular space, and an oligomeric tip [20]. Proteins required for effector translocation (PcrV, PopB, and PopD) are secreted through the needle, and PopB and PopD insert into the host membrane to assemble a hetero-oligomeric transmembrane protein complex or translocon [21-27]. Translocation of effectors into the host cell disrupts cell function and results in a localized disruption of neutrophils [12, 28-31]. Inhibition of translocon assembly will prevent effector translocation and will protect host cells (mainly neutrophils) from the cytotoxic effect of bacterial effector proteins and reduce the virulence of the infecting *P. aeruginosa* strain. Further, since the translocon is assembled and is located outside the bacterial cell [26, 32], small molecule inhibitors would avoid the challenges associated with penetrating the impermeable outer membrane [33] and extrusion by the highly active efflux pumps of *P. aeruginosa* [34, 35].

**Figure 1.**
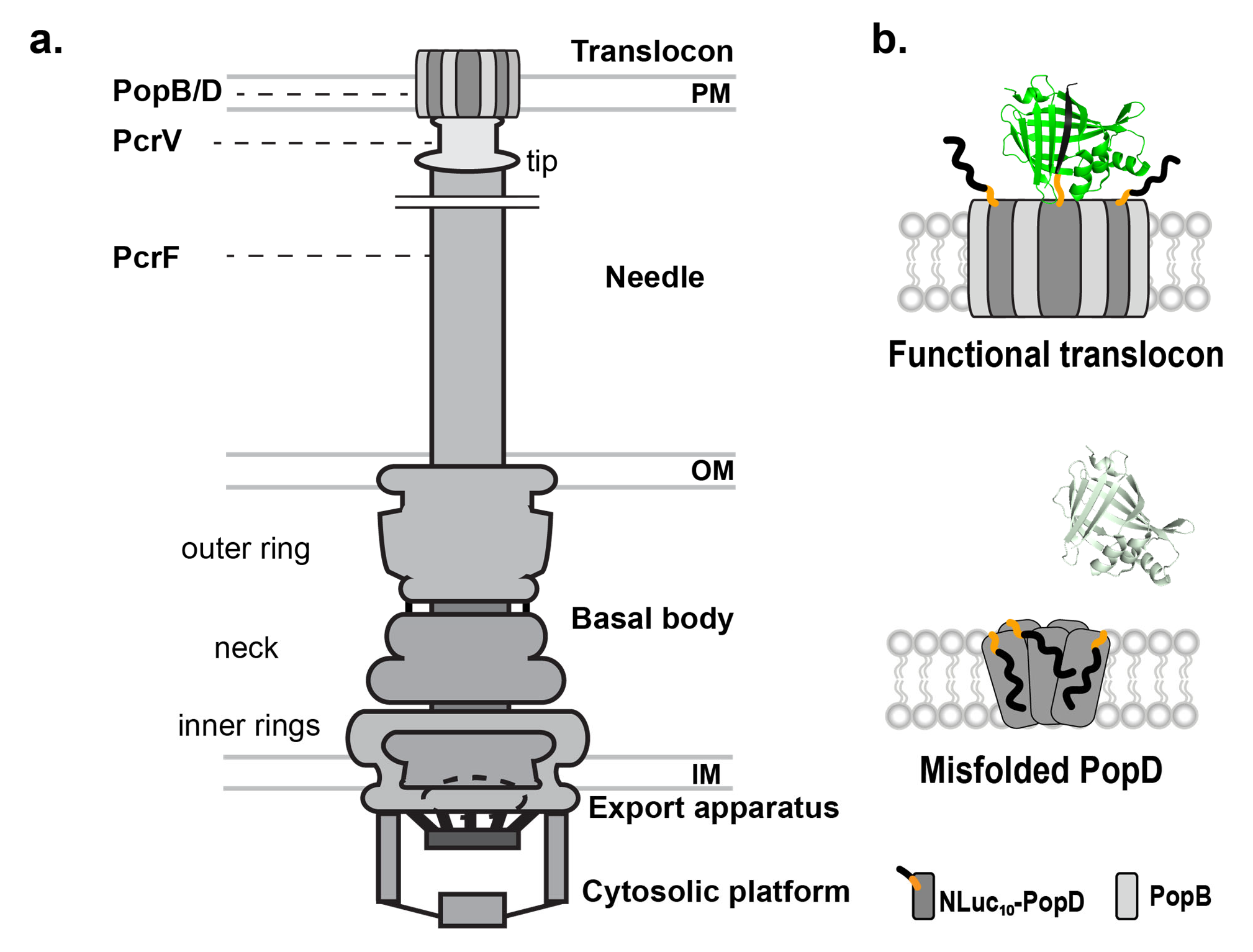
a) Schematic overall structure of a T3SS. The overall apparatus is divided into five structural groups, indicated on the right. The Basal body is formed by three distinct components, indicated on the left. Bacterial membranes and the target membranes, through which secreted protein effectors are translocated are indicated, inner membrane (IM), outer membrane (OM), and plasma membrane (PM). The proteins that form the needle (PcrF), the needle tip (PcrV) and the translocon (PopD and PopB) are indicated. **b) Illustration of Split-NanoLuc assay.** NLuc_1-9_ is expressed in the cell cytoplasm while the NLuc_10_ fragment is fused to the N-terminus of PopD. When PopD is properly inserted, the NLuc_10_ will be exposed to the cell cytosol along with the N- terminus of PopD and will refold with NLuc_1-9_ to form full length NanoLuc (green). Luminescence will be detected only for functional translocons.

In *P. aeruginosa*, the effector translocation process involves the proteins PcrV, PopB, and PopD, which are also referred to as SctA, SctE, and SctB, respectively, following the unified Sct nomenclature, [17, 36]. PcrV is located at the tip of the needle [37], whereas PopB and PopD insert into the host cell membrane [21, 22, 25]. PopB and PopD assemble in an 8:8 ratio to form a 16-member hetero-oligomeric transmembrane complex *in vitro* [24]. PopB is the larger of the translocators and possesses two non-polar segments in its primary structure, while PopD possesses only one non-polar segment [22, 38]. Both proteins adopt a molten globular conformation and are prone to aggregation in aqueous solution [23, 39, 40]. PopB has been shown to assist with the proper insertion of PopD by causing a conformational change, leaving the N-terminal region of PopD exposed to the host cytosol only when forming functional translocons [41]. Hence, the molecular details revealed by the *in vitro* and *in vivo* studies of the translocators provided the basis to develop an assay that reports on proper translocon assembly. This novel assay can be used to identify inhibitors of PopD assembly into functional translocons.

In this study, we developed a whole-cell infection assay that detects inhibitors of T3SS translocon assembly in *P. aeruginosa*. With the distinct conformation adopted by PopD in functional translocons, a split NanoLuc™ luciferase system [42] was developed to identify compounds that interfere with the translocon assembly. Here, we report the construction and optimization of a whole-cell-based assay, and the analysis of a pilot screen using an FDA-approved compound library. Primary hits were assessed using secondary and ortholog assays, developing a workflow to prioritize hit compounds readily. These results showed that the assay is robust and appropriate for the expansion to a high-throughput screen for novel translocon assembly inhibitors. Moreover, the assay provides a signal for functional translocon assembly that could be exploited to study the structure and assembly of T3SS translocons on cellular membranes in *P. aeruginosa* and related pathogens.

## Results and Discussion

### Experimental rationale

In a previous study we demonstrated that the N-terminus of PopD is exposed to the cytosol of the target-cell only when PopD and PopB are assembled in functional translocons [41]. Infection of HeLa cells with a *P. aeruginosa* strain lacking PopB, which is unable to assemble a functional translocon, showed that PopD was stably inserted into the host membrane, but its N-terminus was not exposed to the cytosol of the target cell [41]. We hypothesized that by taking advantage of the distinct exposure of the PopD N-terminus to the cytosol of target cells, we could develop an assay to effectively monitor the assembly of the translocon. To achieve this objective, we employed the split NanoLuc protein complementation reporter system [42], which has demonstrated success in studying diverse intermolecular interactions, including the secretion of T3SS effectors [43, 44]. The rationale for the assay is illustrated in Figure 1b. The NanoLuc system is comprised of a stability optimized N-terminal 18 kDa fragment containing β-strands 1-9 (NLuc_1-9_) and an affinity-optimized 1.3 kDa C-terminal fragment or β- strand 10 (NLuc_10_). Neither fragment alone produces a significant luminescence signal when incubated with the substrate [42], however, when NLuc_10_ binds to NLuc_1-9_, it assembles into an active enzyme. To utilize this system for monitoring the conformation of PopD, we constructed a strain of *P. aeruginosa* PAK that expresses an N-terminal NLuc_10_-PopD fusion protein and a HeLa cell line that constitutively expresses NLuc_1-9_, as described below. Thus, proper insertion and assembly of the T3SS translocon will result in reconstitution of NanoLuc in the cytoplasm of the target cell that will generate a bioluminescent signal in the presence of substrate. Conversely, inhibition of translocon assembly would prevent NanoLuc complementation, resulting in a loss of signal. Hence, this assay system holds the potential to investigate the assembly of the *P. aeruginosa* translocon and identify inhibitors that can target this crucial process.

### Generating the *P. aeruginosa* PAK strain producing NLuc_10_-PopD fusion protein

Detecting the proper assembly of the *P. aeruginosa* T3SS translocon required a PAK strain that expresses an NLuc_10_-PopD fusion protein. The DNA fragments encoding NLuc_10_ and a flexible peptide linker (GSSGGSSG) were assembled into the plasmid pUCP18-GHD to generate the pNLuc_10_-popD plasmid as described in Experimental Section (Figures 2a and S1a). As shown in Figure 2b, the NLuc_10_-PopD fusion protein supports translocation of the ExoS effector (encoded in PAKΔpopD strain) into HeLa cells, as evidenced by the rounding of infected cells [24]. The generated pNLuc_10_-popD plasmid was then transformed into PAKΔexsEΔexoSTYΔpopD (hereafter PAKΔEΔSTYΔD) strain [24], which carries the following genetic modifications to improve performance in the assay: a) deletion of the exsE gene increases expression of the T3SS regulon [45]; b) deletion of genes encoding the ExoS, ExoT, and ExoY effector proteins minimizes target cell damage after infection [41]. In addition to the PAKΔEΔSTYΔD(pNLuc_10_-popD) strain that produces a functional translocon, the PAKΔEΔSTYΔDΔpopB(pNLuc_10_-popD) strain was used as a positive control for inhibition, given that PopD inserts into the membrane in the absence of PopB, but its N-terminus is not exposed to the cytosol [41].

**Figure 2.**
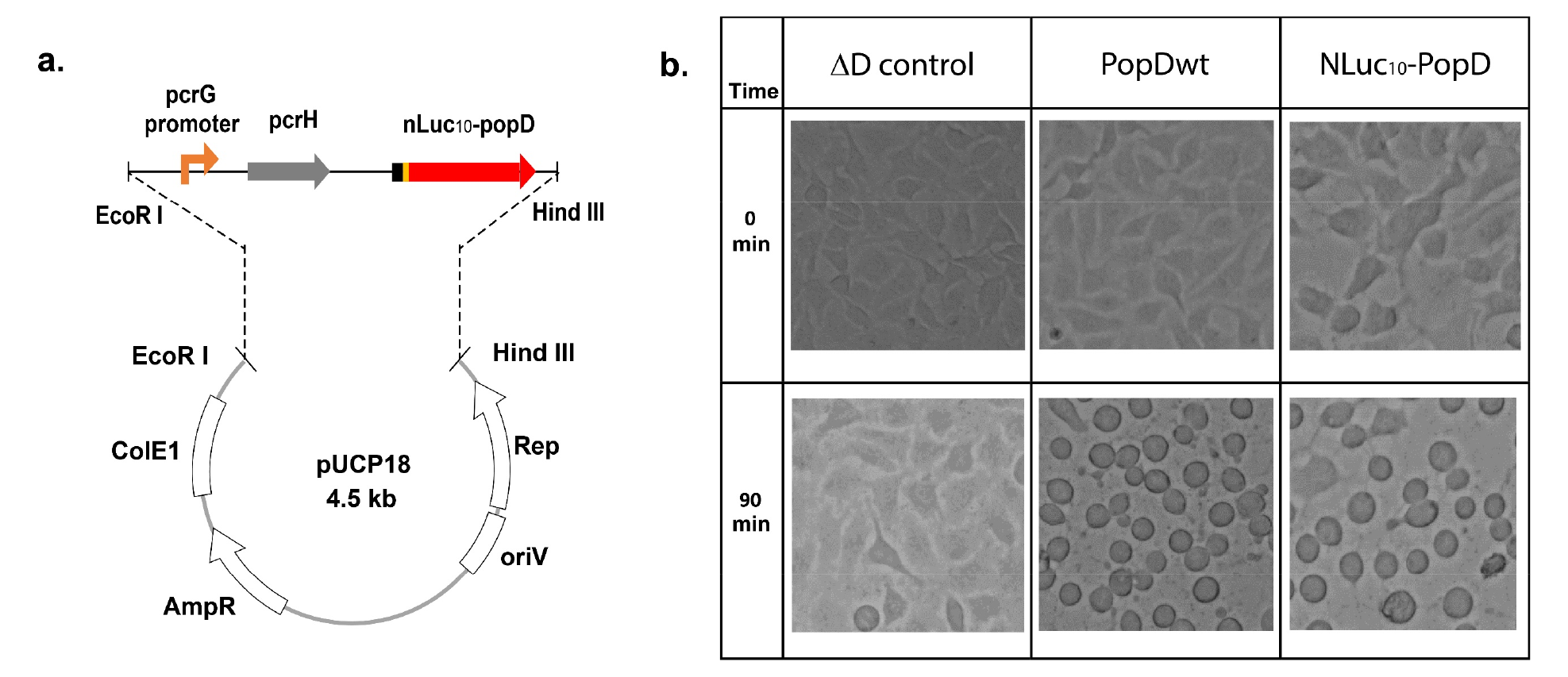
Construction of pNLuc_10_-popD plasmid. **a) Scheme of the plasmid used to express NLuc_10_-PopD in PAK strains.** The intrinsic promoter regulates the expression of the NLuc_10_-PopD and cognate chaperone PcrH. The target fragment was inserted between EcoR I site and Hind III site on the *Escherichia*-*Pseudomonas* shuttle vector pUCP18. The plasmid contains high-copy-number ColE1 origin of replication and broad-host-range origin of replication, oriV, which comes from *P. aeruginosa*. The plasmid also contains Ampicillin resistant gene (AmpR) for selection and replication protein gene (Rep) from *P. aeruginosa*. **b) Functionality of NLuc_10_-PopD using the rounding assay**. HeLa cells were infected with PAKΔD(pUCP18), PAKΔD(pUCP18-GHD) or PAKΔD(pNLuc_10_-popD) at MOI = 30. Expressed PopD variant or control without PopD is indicated at the top. Time of infection is indicated at the left. Images were taken by an inverse microscope.

### A stable HeLa cell line expressing NLuc_1-9_ provides a robust platform for the assay

A stable cell line that constitutively expresses NLuc_1-9_ was created to provide uniform NLuc_1-9_ expression levels when the assay is performed in multi-well plates. HeLa cells were chosen for the infection assay because they were more adherent and resistant to detachment than other commonly used cell types (e.g., HEK293), allowing for the multiple washes post infection and long incubation required for the complementation of the NanoLuc fragments after bacterial infection. Consequently, a stable HeLa cell line expressing NLuc_1-9_ was created as described in the Experimental Section. Briefly, the coding region for NLuc_1-9_ was cloned into a lentiviral vector, resulting in pLenti-NLuc_1-9__IRES_EGFP, which expresses NLuc_1-9_ under a CMV promotor and co-express EGFP by using the internal ribosome entry site 2 (IRES2) sequence. HeLa cells were transfected with lentiviral particles containing pLenti-NLuc_1-9__IRES and stable integrants were selected with 1 mg/ml G418, evaluating GFP expression by flow cytometry. The final cell line, named NLuc_1-9_-HeLa, exhibited more than 200-fold increase in EGFP expression as compared to the parent strain.

### Initial assay optimization using manual conditions

With the PAKΔEΔSTYΔD(pNLuc_10_-popD) strain and the NLuc_1-9_-HeLa cell line in hand, the next step was the optimization of the assay to monitor the inhibition of translocon assembly. In brief, NLuc_1-9_-HeLa cells were infected with the PAKΔEΔSTYΔD(pNLuc_10_-popD) strain, and after removal of bacteria, cells were further incubated to allow the assembly of the complementary NanoLuc fragments (Figure 3a). An MOI of 50 was used based on the observed maximum insertion of translocators and minimum detachment of HeLa cells [46]. The number of HeLa cells per well was kept at ∼10,000. Various infection and complementation times were evaluated to select optimal conditions for the assay and the results are shown in Figures 3b and 3c. Half maximal signal was observed after 15 min of infection, with maximal signal reached after 45 min (Figure 3b). These results suggest that the maximal number of NLuc_10_ fragments exposed to the target cell’s cytosol reached saturation, and that this saturation was independent of any effect of the ExoY, ExoS, or ExoT effectors into the host cell. Using an optimal infection time of 45 min, we determined the effect of additional complementation time on the overall luminescent signal. As shown in Figure 3c, the maximal signal was achieved after 30 min of incubation. Any effect on the overall signal resulting from the degradation of NLuc_10_ post-infection is unlikely because the signal increase continued during complementation, after washing out the bacteria. In summary, maximal signal was reached after 45 min of infection, followed by 30 min of additional complementation and these experimental conditions were adopted for the assay.

**Figure 3.**
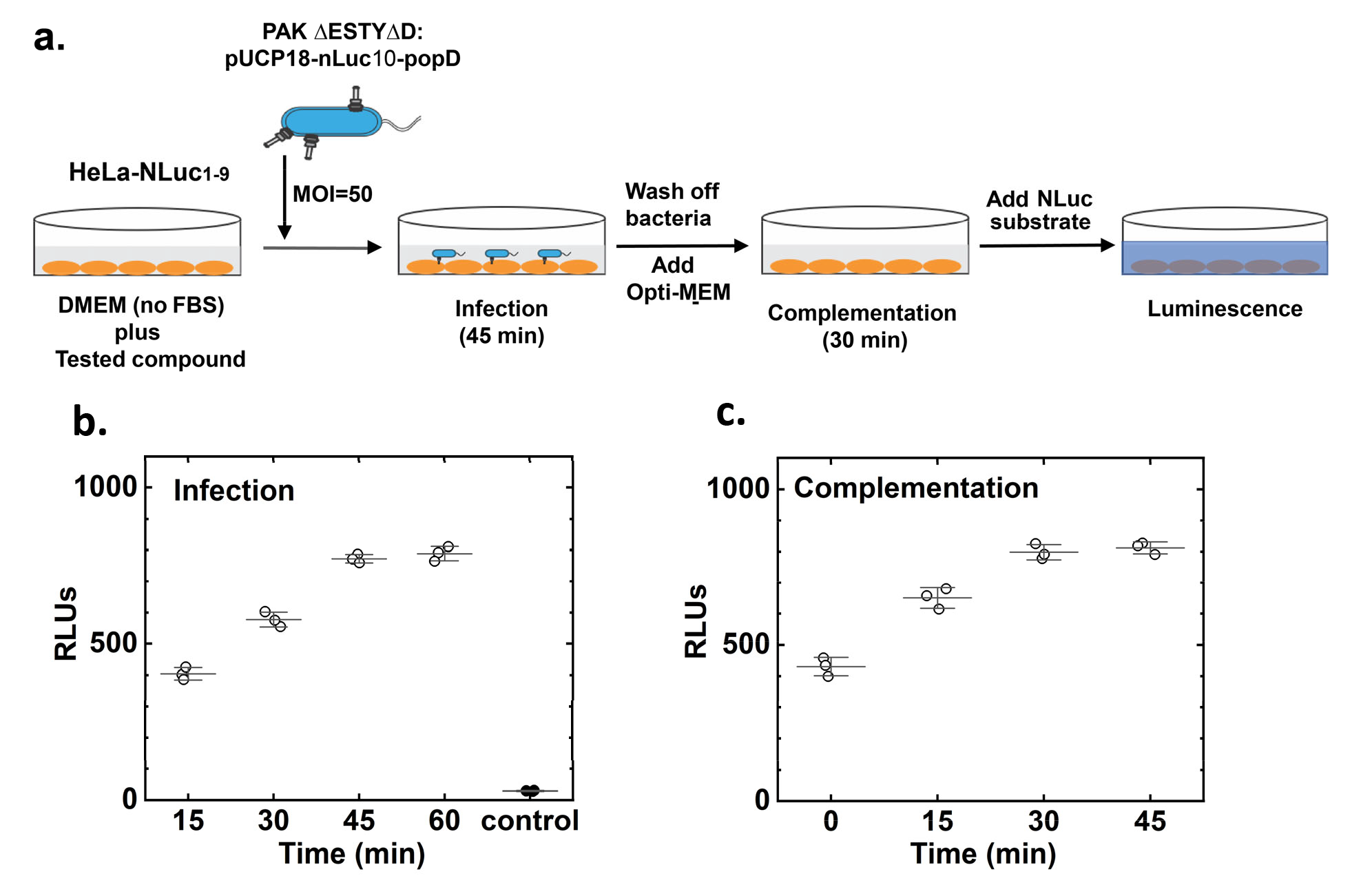
Scheme of the assay procedure and optimization. **a)** Bacteria were added to the ∼10000 cells at an MOI of 50 in the presence of the compound to be tested and incubated for 45 min (infection period). After infection the bacteria were washed off and the culture media changed to Opti-MEM to eliminate Phenol red that interferes with the luminescent signal. Cells are incubated for an additional 30 min to maximize NanoLuc complementation. The NanoLuc reagent is added, and the luminescence measured in a plate reader. **b)** NLuc_1-9_-HeLa cells were incubated with the strain PAKΔEΔSTYΔD(pNLuc_10_-popD) for the indicated times (open circles). Control without bacteria was measured after 60 min (filled circles). Complementation time was fixed at 30 min. **c)** NLuc_1-9_-HeLa cells were incubated with the strain PAKΔEΔSTYΔD(pNLuc_10_-popD) for 45 min and additionally incubated for the indicated times after washing off bacteria. Each long-centered line shows the average of three replicas and the error bars represent the standard deviation.

Next, the variability of the assay was evaluated using experimental replicas for the positive and the negative inhibition control (Figure 4a). As described above, we used the PAKΔEΔSTYΔDΔpopB(pNLuc_10_- popD] strain as a control for positive inhibition of translocon assembly, and the PAKΔEΔSTYΔD(pNLuc_10_-popD) strain as a negative control of inhibition. After incubation with the PAKΔEΔSTYΔD(pNLuc_10_-popD) strain, the signal produced by NLuc_1-9_-HeLa cells (555 ± 23) was 10 times higher as compared to the background signal (55 ± 2) from the non-infected cells (Figure 4a). When using the PAKΔEΔSTYΔDΔpopB(pNLuc_10_-popD) strain as a control for inhibited translocon assembly, the relative luminescence signal was 135 ±14, approximately 75% lower than that of the non-inhibited signal [note that in the absence of PopB, PopD is still inserted into the membrane but in a non-functional conformation 41]. The signal to noise ratio (S/N) for the positive and negative inhibition controls was 17.9. To determine whether the assay is robust, we calculated the Z’ factor using data obtained from nine technical replicates each of the positive inhibition control and the negative inhibition control and a Z’ factor of 0.73 was obtained, indicating that the assay is suitable for screening of translocon inhibitors. The Z’ factor is a coefficient defined to reflect both the assay signal dynamic range and the data variation associated with the signal measurements. An ideal assay will produce a Z’ factor score of 1 and any assay that has 1> Z’ ≥0.5 can be considered as an excellent assay with a large separation band [47].

**Figure 4.**
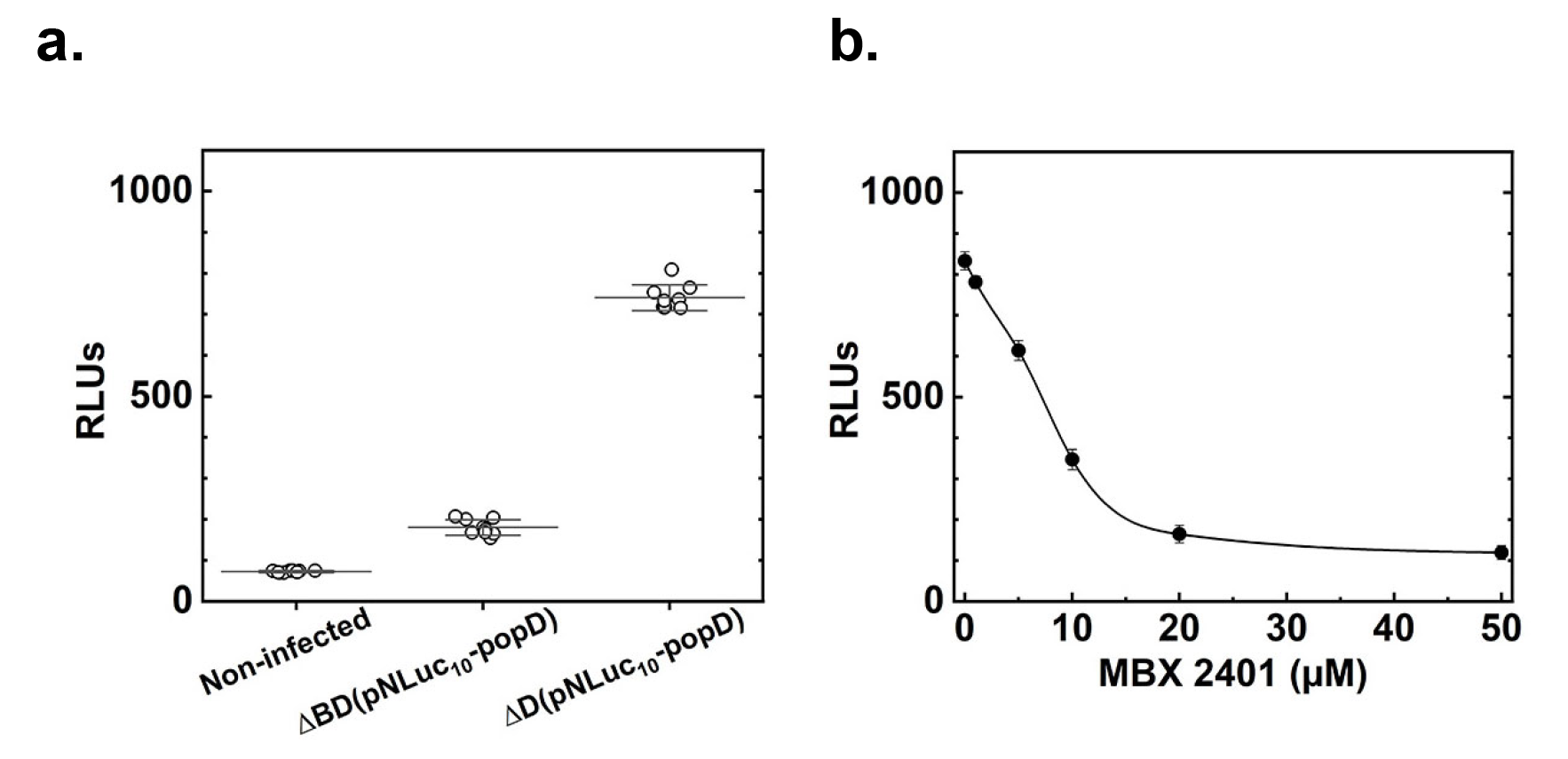
Split NanoLuc assay is excellent for screening and sensitive to inhibitor concentrations. **a)** Each group of data correspond to Non-infected NLuc_1-9_-HeLa cells (left); NLuc_1-9_-HeLa cells infected with strain PAKΔEΔSTYΔDΔpopB(pNLuc_10_-popD) (center); and NLuc_1-9_-HeLa cells infected with strain PAKΔEΔSTYΔD(pNLuc_10_- popD) (right). Long line shows the average of nine replicas. **b)** NLuc_1-9_-HeLa cells were incubated with the strain PAKΔEΔSTYΔD(pNLuc_10_-popD) in the presence of 0.5% DMSO or the indicated concentrations of MBX-2401 in

To validate the sensitivity of the assay to small-molecule inhibitors, we measured the half-maximal inhibitor concentration (IC_50_) of MBX-2401, a (R)-isomer of a phenoxyacetamide T3SS inhibitor [48], in the cell-based translocon assembly assay. The response of the assay to varying concentrations of MBX- 2401 is shown in Figure 4b. The calculated IC_50_ was 8 µM, which is in good agreement with previously reported values [16, 48]. This result indicated that the PopD translocon assembly assay is sensitive to small molecule inhibitors of the T3SS. However, it is likely that MBX-2401 is an indirect inhibitor of translocon assembly, as it inhibits both secretion and translocation of proteins that are extruded through the needle complex, including PcrV, PopD, and PopB [48]. Therefore, it will be essential to include a secondary screen for T3SS secretion inhibitors into the workflow of the screening procedure to distinguish translocon assembly inhibitors from those inhibitors affecting downstream events like transcription, translation, or secretion of T3SS components.

### Similar optimal parameters were required for automated high throughput screening conditions

To adapt the PopD assembly assay to a high throughput format suitable for screening compound libraries, standard laboratory automation was utilized for the transfer of reagents and for washing steps, as described in the Experimental Section. During the implementation of automation equipment into the assay workflow, important assay variables including multiplicity of infection (MOI), bacterial infection time, and complementation time were re-evaluated in 96-well plates. It was observed that the optimal conditions reported for the manual assay (MOI 50, 45 min for infection, and 30 min for NLuc complementation) remained the optimal conditions for the 96-well plate assay (data not shown). However, instead of using a PAKΔEΔSTYΔDΔpopB(pNLuc_10_-popD) strain as the positive control of inhibition for the automated system, it was more practical to use the small-molecule inhibitor of T3SS secretion (MBX-2401) as positive control and homogenously deliver a single PAK strain into all wells.

We repeated the procedure using three identical 96-well plates, on two different days, to assess the robustness of the automated PopD assembly assay. Half of the wells in each plate contained 1% DMSO (negative control for inhibition) and the other half were treated with 10 µM MBX-2401 (positive control for inhibition). The data from 6 plates were consistent, with an average percent coefficient of variation (Standard Deviation/Mean x 100) of 17% and 13% for the negative control and positive control, respectively, an average Z’ of 0.23, and an average S/N ratio of 4.6. As expected for the automated assay, the average Z’ and the S/N ratio were lower than that observed for the manual assay used during assay development. This is in part due to the automatic washes, that are less efficient and harder on attached cells than the washes performed manually. Given the complexity of the assay and the small distribution observed for positive control samples, we considered a Z’ of 0.23 sufficient to discriminate whether a compound inhibits the assembly of PopD [47].

### Performance and reproducibility of the assay when testing a small-compound library

A commercially available collection of 776 discrete chemical compounds comprised of FDA-approved drugs was used to evaluate the performance of the assay in a small-scale pilot screen, using the conditions optimized above. Each assay plate contained negative and positive inhibition controls in columns 1 and 12, respectively. Library compounds dissolved in DMSO were added to each well in columns 2-11 at a final concentration of 25 µM (final DMSO concentration was 1%) using a Caliper SciClone ALH 3000 liquid handler equipped with a 96-tip pipetting head.

Two replicates of the FDA library screen were performed on different days to assess the reproducibility of the assay (Figure 5a and S2). The variability in luminescence signal for the positive (green) and the negative (red) inhibition controls used in the screen was analyzed using a box plot and a normal distribution, and compared with the data obtained when samples were treated with each library compound. The first replica showed that the average signal for the negative inhibition control (101856 ± 22259) decreased 8.3 times in the positive inhibition control samples (12320 ± 2259). The Z’ score for this data set was 0.34. For the second replica, the average signal for the negative inhibition control (111826 ± 26723) was reduced 5.4 times in the positive inhibition control samples (20685 ± 6888). The Z’ score for this data set was −0.11, indicating some overlap between the separation bands. However, a close inspection of the data distribution showed that a consistent set of low number of hits was obtained in both replicas (Figure S2). The variability in the Z’ between replicas was mostly due to the dispersion of the negative inhibition control values, which are more sensitive to the number of cells per well and the number of added bacteria. This variability is not surprising since the manipulation of samples is less uniform when using automated sample dispensers. However, when inhibition occurs, the lack of signal is not significantly influenced by the variability on the number of cells used in the assay.

**Figure 5.**
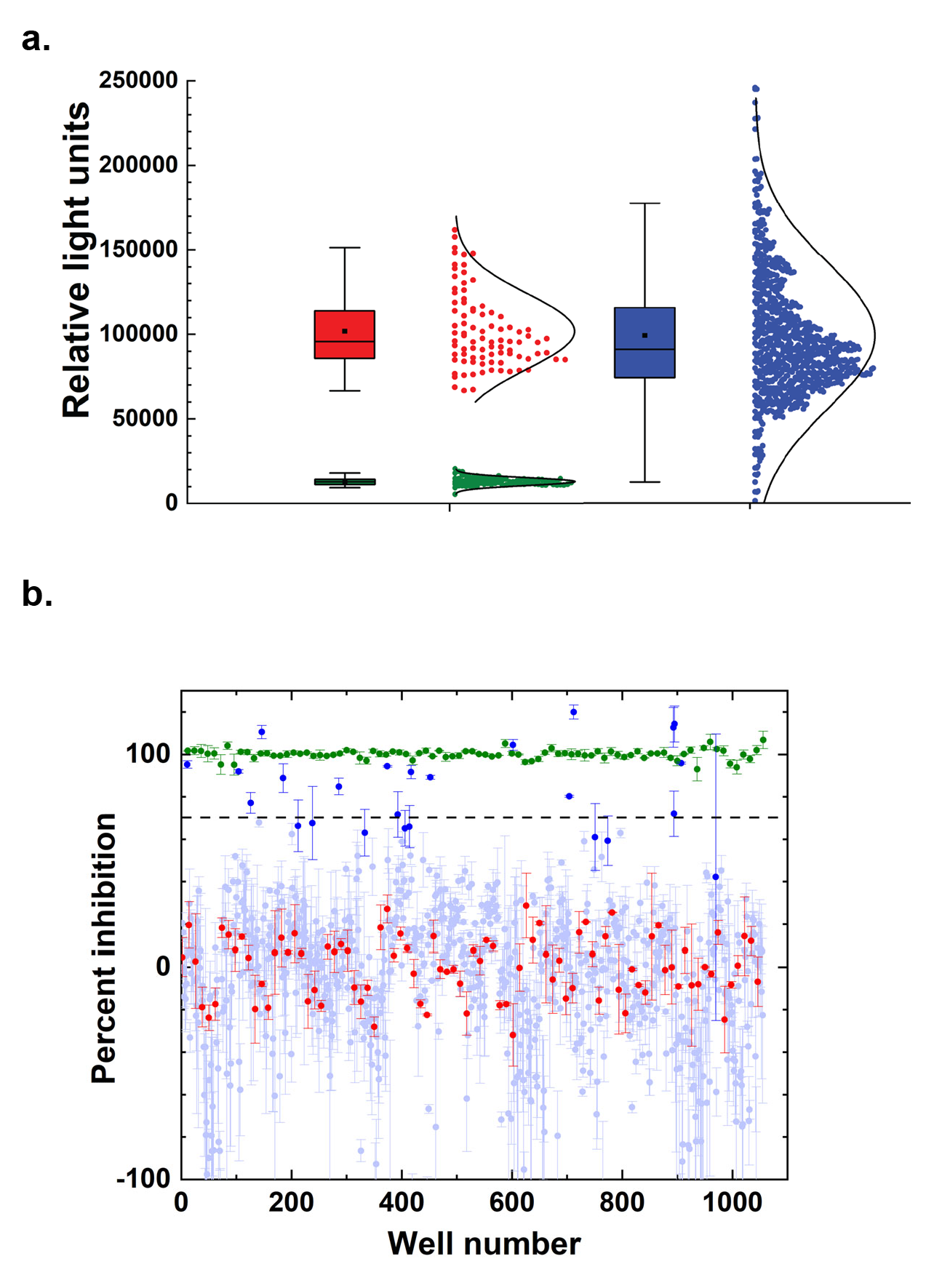
Variability of the assay for the screening of the FDA compound library. **a)** Raw data in relative light units were plotted as a box plot with normal distribution. The negative control (DMSO-treated) is shown in red, the positive control (MBX-2401) is shown in green, and screened compounds are shown in blue. The average S/B was 8.3, with an average Z’ value of 0.34 across eleven 96 well plates. The box plot shows the mean (black square), the median (line), the 25-75 percentage range (boxes), and the standard deviation (whisker). **b)** Percent of inhibition for the positive controls (green), negative controls (red), and screened compounds (blue). Filled circles correspond to the average of the two replicas and the error bars to their range. The horizontal dashed line shows the threshold used to select hits (70 % inhibition). Selected hits are shown in dark blue.

A cutoff to select positive hits in the assay was determined using normalized data to compare the results obtained on different data sets. The raw data from each plate was normalized by calculating the percentage of inhibition for each compound using the negative and positive inhibition control values as described in the Experimental Section. The number of hits obtained on each replica was plotted as a function of the selected threshold (Figure S3). For this small library, a cutoff of ≥70% inhibition was selected to set the number of hits lower than ∼3% of all screened compounds. Using this cutoff, we identified 24 hits from replica 1 (hit rate: 3%), and 16 hits from replica 2 (hit rate: 2%), with a total of 25 unique hits (primary hit rate 3.2%) from the two replicates (Figure 5b). To confirm the activity of the 25 primary hits, the compounds were re-tested in the PopD assembly assay in triplicate, and 14 of 25 primary hits exhibited an average of ≥70% inhibition in the confirmation assay (confirmed hit rate 1.8%, Table 1).

**Table 1.**
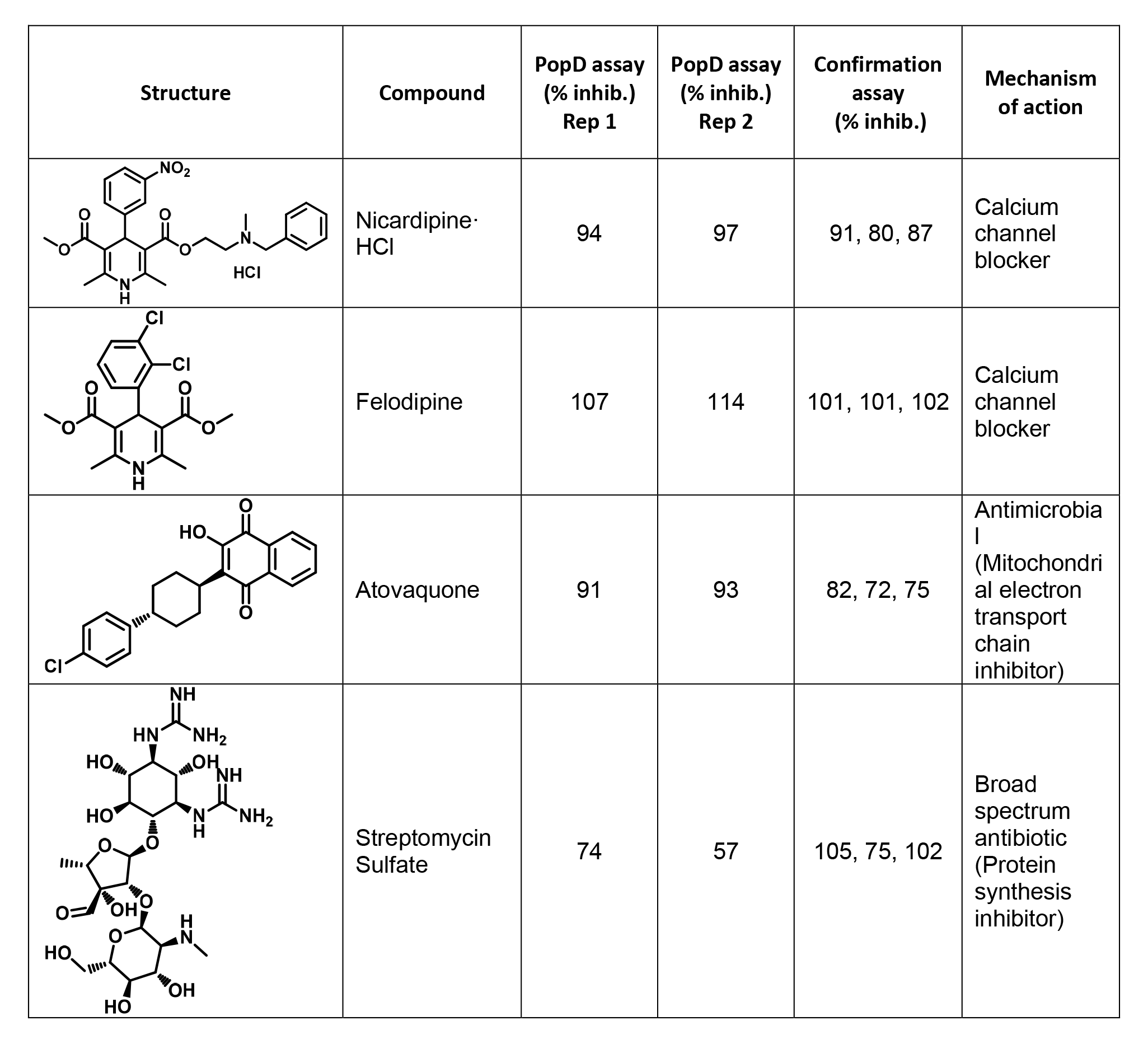
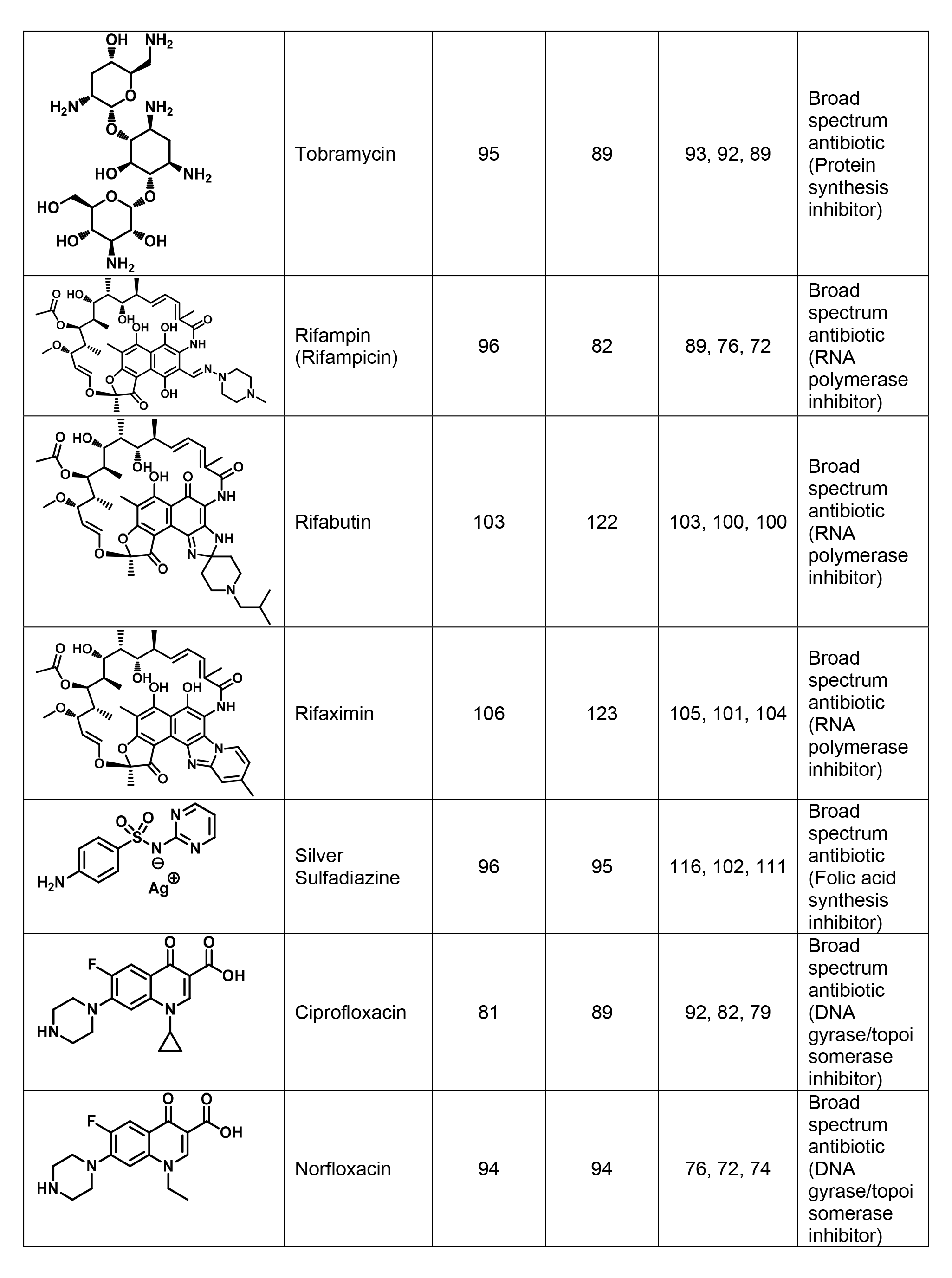
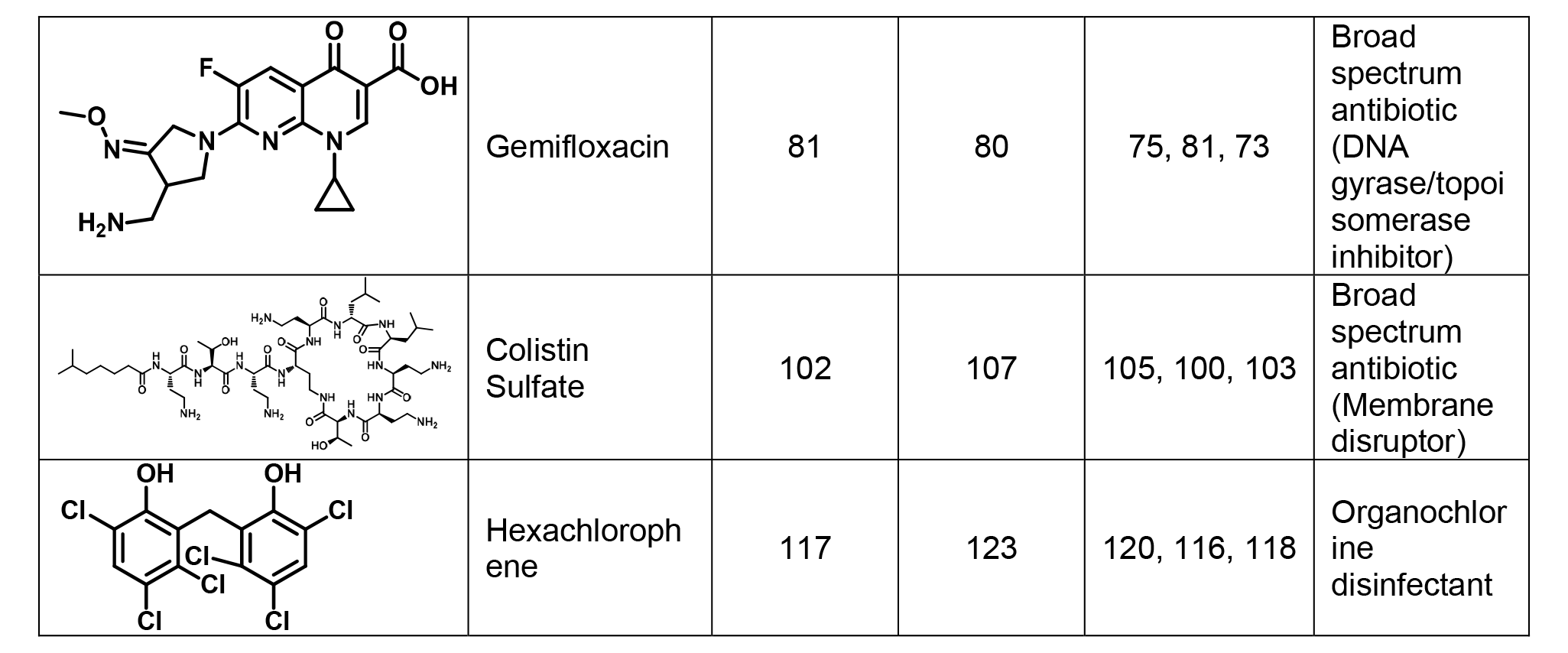
Confirmed hits from the PopD assembly assay, with compound structure, name, and mechanism of action reported along with % Inhibition calculated from the two runs of the assay and from hit-confirmation studies. Any compound that reached ≥70% inhibition in the PopD assembly assay in at least one replicate of screening was cherry-picked and tested in a hit conformation assay. All compounds were evaluated at 25 µM. Confirmation assays were performed in technical triplicate.

| Structure | Compound | PopD assay<br>(% inhib.)<br>Rep 1 | PopD assay<br>(% inhib.)<br>Rep 2 | Confirmation<br>assay<br>(% inhib.) | Mechanism<br>of action |
| --- | --- | --- | --- | --- | --- |
|  | Nicardipine-HCl | 94 | 97 | 91, 80, 87 | Calcium channel blocker |
|  | Felodipine | 107 | 114 | 101, 101, 102 | Calcium channel blocker |
|  | Atovaquone | 91 | 93 | 82, 72, 75 | Antimicrobial<br>(Mitochondrial electron transport chain inhibitor) |
|  | Streptomycin Sulfate | 74 | 57 | 105, 75, 102 | Broad spectrum antibiotic<br>(Protein synthesis inhibitor) |
|  | Tobramycin | 95 | 89 | 93, 92, 89 | Broad spectrum antibiotic (Protein synthesis inhibitor) |
|  | Rifampin (Rifampicin) | 96 | 82 | 89, 76, 72 | Broad spectrum antibiotic (RNA polymerase inhibitor) |
|  | Rifabutin | 103 | 122 | 103, 100, 100 | Broad spectrum antibiotic (RNA polymerase inhibitor) |
|  | Rifaximin | 106 | 123 | 105, 101, 104 | Broad spectrum antibiotic (RNA polymerase inhibitor) |
|  | Silver Sulfadiazine | 96 | 95 | 116, 102, 111 | Broad spectrum antibiotic (Folic acid synthesis inhibitor) |
|  | Ciprofloxacin | 81 | 89 | 92, 82, 79 | Broad spectrum antibiotic (DNA gyrase/topoisomerase inhibitor) |
|  | Norfloxacin | 94 | 94 | 76, 72, 74 | Broad spectrum antibiotic (DNA gyrase/topoisomerase inhibitor) |
|  | Gemifloxacin | 81 | 80 | 75, 81, 73 | Broad spectrum antibiotic (DNA gyrase/topoisomerase inhibitor) |
|  | Colistin Sulfate | 102 | 107 | 105, 100, 103 | Broad spectrum antibiotic (Membrane disruptor) |
|  | Hexachlorophene | 117 | 123 | 120, 116, 118 | Organochlorine disinfectant |

The FDA-approved compound library included numerous bactericidal and bacteriostatic compounds. Hence, further secondary assays are required to distinguish between compounds that inhibit the secretion of translocators and those that impede the assembly of a fully functional translocon.

### Overall strategy to evaluate and confirm T3SS translocon inhibitors

The injection of the T3SS effectors into the target cells is a multi-step process, that involves gene transcription, mRNA translation, protein secretion, interaction with the host, and ends with translocation of proteins across the plasma membrane of the host cell. The assembly of the translocon complex is a late step in this process. A positive hit in the PopD assembly assay will indicate that the translocon was not assembled properly. However, compounds perturbing earlier steps of the injection process, having bactericidal properties, or cytotoxicity for the target cell, could also be detected as positive hits in the assay. To specifically identify inhibitors that hinder the assembly of PopD into functional translocons, we devised a strategy to discriminated true hits from compounds affecting other T3SS processes or processes unrelated to the functionality of the T3SS (Figure 6). We used this strategy to evaluate the confirmed hits identified in the FDA library during the pilot screen.

**Figure 6.**
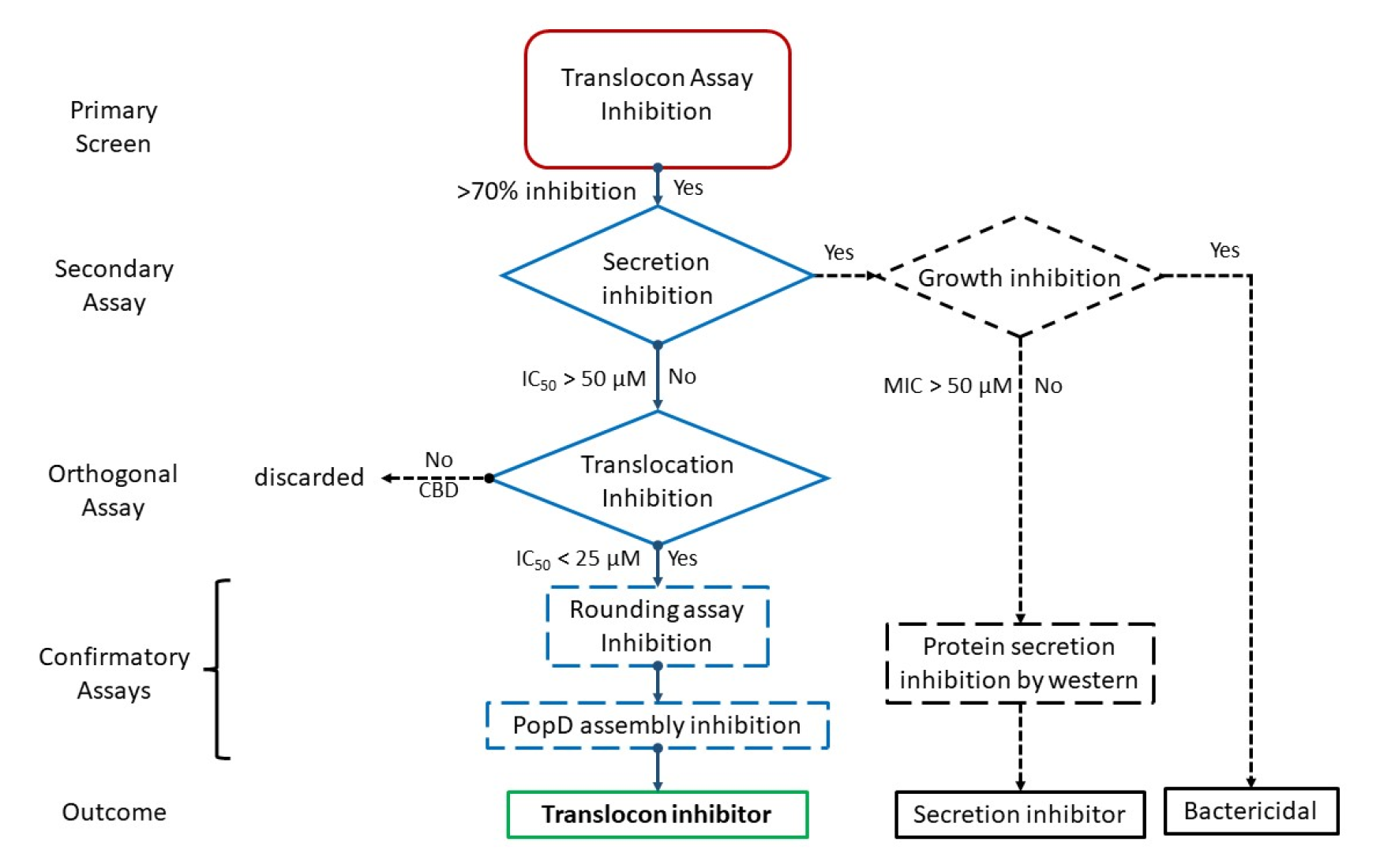
Flow chart of the screening strategy. Positive hits from the primary screen (red box) are evaluated for their effect on protein secretion as described in the Experimental Section. Those that do not do not block secretion are evaluated for translocation inhibition using an ortholog assay. Finally, translocon inhibitors are confirmed using the rounding assay on a PAK wild type strain, and for their specific activity on PopD assembly. Compounds that inhibited bacterial growth are deprioritized and classified as bactericidal. Those compounds that inhibited secretion, but did not inhibit bacterial growth are deprioritized, and could be confirmed as secretion inhibitors using western blot. CBD cannot be determined.

Following the primary screening, compounds exhibiting more than 70% inhibition will undergo triplicate re-testing. Confirmed hits will then be evaluated for their ability to inhibit secretion. Compounds inhibiting secretion with an IC_50_ ≤ 50 µM will be tested for bacterial growth inhibition. Those compounds with an MIC < 25 µM are categorized as bactericidal or bacteriostatic. Compounds with an MIC > 50 µM will be deprioritized and classified as potential T3SS secretion inhibitors. Inhibition of secretion could be confirmed through western blot detection of secreted proteins, including translocators and effectors. It is important to test the secretion of both translocators and effectors as their secretion is regulated by distinct mechanisms [49-51].

Compounds that did not inhibit secretion will undergo evaluation for translocation inhibition using an orthogonal assay, to account for any artifact caused by inhibitors on the NanoLuc enzyme. Compounds that fail to inhibit translocation-dependent HeLa cell lysis or exhibit toxicity towards HeLa cells will be discarded. Finally, compounds that inhibit the orthogonal assay will be confirmed by their ability to inhibit translocation using the morphological changes observed in HeLa cell infected by PAK wild type strain [rounding assay, 52]. Furthermore, the specificity of inhibition on PopD assembly, particularly N-terminus exposure, will be verified through western blot analysis using the anti-phospho-GSK antibody, as described previously [46].

In conclusion, the comprehensive strategy incorporates multiple assays to identify and categorize inhibitors of the T3SS. Through a series of assays, we aim to differentiate true hits from artifacts, evaluate the presence of other antibacterial compounds not acting on the T3SS, assess the presence of inhibitors that affect T3SS secretion, and select for those compounds that specifically block protein translocation. This rigorous approach ensures the thorough evaluation of compound libraries and provides a robust framework for the discovery of novel T3SS inhibitors.

### Evaluation of the procedure to confirm and classify positive hits

The minimum inhibitory concentrations [MICs, 53] of the fourteen hit compounds were determined against *P. aeruginosa* PAKΔEΔSTYΔD(pNLuc_10_-popD), the strain used in the primary assay. Among the fourteen compounds, ten exhibited MIC values ≤ 50 µM, which aligns with the presence of various broad-spectrum antibiotics in this group of hits (Table 1 and 2).

Subsequently, compounds that did not significantly affected bacteria growth were subjected to analysis for T3SS protein secretion using the luciferase transcriptional reporter, as described in the experimental section [54]. This assay measures activation of the T3SS promotors triggered by the binding of the master regulator ExsA, which is released through the secretion of the T3SS regulator ExsE [45]. None of the four compounds showed T3SS secretion inhibition (Table 2).

**Table 2.** Evaluation of confirmed hits from the PopD assembly assay using secondary and orthogonal assays. Compounds that inhibited growth (MIC) or secretion were de-prioritized. Compounds were subsequently assessed for T3SS protein translocation and compounds that failed to inhibit cell lysis were discarded. Assays were performed in technical triplicates. n.d. = not determined.

| Compound | Inhibition<br>Primary<br>assay (%) | MIC<br>(μM) | Secretion<br>IC<sub>50</sub> (μM) | Translocation<br>IC<sub>50</sub> (μM) |
| --- | --- | --- | --- | --- |
| Nicardipine-HCl | 95 | >50 | >50 | >100 |
| Felodipine | 110 | >50 | >50 | CBD |
| Atovaquone | 92 | >50 | >50 | >100 |
| Streptomycin<br>Sulfate | 74 | >50 | >50 | >100 |
| Rifampin<br>(Rifampicin) | 89 | 50 | 22.1 ± 1.5 | n.d. |
| Rifabutin | 113 | 25 | 9.1 ± 0.7 | n.d. |
| Hexachlorophene | 120 | 25 | 6.3 ± 0.1 | n.d. |
| Rifaximin | 115 | 13 | 2.4 ± 0.2 | n.d. |
| Silver Sulfadiazine | 96 | 13 | 0.7 ± 0.1 | n.d. |
| Ciprofloxacin | 85 | 6.3 | 1.0 ± 0.1 | n.d. |
| Colistin Sulfate | 105 | 3.1 | <0.098 | n.d. |
| Tobramycin | 92 | 3.1 | 2.3 ± 0.1 | n.d. |
| Norfloxacin | 94 | 1.6 | 22 ± 12 | n.d. |
| Gemifloxacin | 80 | 1.6 | 1.1 ± 0.1 | n.d. |

Confirmed hits that did not inhibit secretion were further evaluated for effector translocation inhibition using an orthogonal assay (Figure 6), which relies on the T3SS-dependent death of CHO cells [55]. Cell death was measured by the release of the cytosolic enzyme lactate dehydrogenase (LDH), produced by the translocation of the *P. aeruginosa* PA99 effector ExoU, a phospholipase that causes CHO cell lysis.

Inhibitors of translocon assembly would prevent effector translocation and consequent cell lysis. However, since a compound itself can be toxic to HeLa cells and induce LDH release, positive hits were evaluated based on their effect on cell viability in the absence of bacteria. Felodipine was found to be toxic to HeLa cells and was discarded. The remining three compounds did not inhibit translocation in the orthogonal assay and were also discarded (Table 2). Significantly, the calcium channel blockers (nicardipine and felodipine) carry a 1,4-dihydropyridine moiety that can also act as inhibitors of NanoLuc [56]. Therefore, it is likely that nicardipine and felodipine were identified as primary hits because they inhibited NLuc reporter activity.

In summary, out of the initial 776-compound library used in this high throughput screening test, 25 compounds were selected as primary hits (Table 3). Following cherry picking and triplicate testing, 14 hits were confirmed. By setting the inhibition threshold at 70%, less than 2% of the compounds were confirmed as hits. While this is a relatively high percentage of positive hits for a high throughput screen, the threshold can be adjusted depending on the size of the screened library. Among these 14 positive hits, 10 compounds exhibited an MIC ≤ 50 µM and were deprioritized. These 10 compounds are widely recognized as broad-spectrum antibiotics or bactericides, and they were satisfactorily classified as non-T3SS inhibitors of PAK in the evaluation procedure. The remaining compounds consisted of two known calcium channel blockers (nicardipine and felodipine), one antimicrobial (atovaquone), and one antibiotic (streptomycin). Due to its toxicity to CHO cells, felodipine was discarded as a potential inhibitor. The other 3 compounds showed no inhibition of translocation when tested using the orthogonal assay.

**Table 3.** Analysis for the PopD assembly assay screening using the FDA compound library.

| Screening stage | Number of<br>compounds | % of<br>Total | Criteria |
| --- | --- | --- | --- |
| Total screened | 776 | 100 | FDA library |
| Primary hits | 25 | 3.2 | Inhibition $\geq 70\%$, 2 independent replicas |
| Confirmed hits | 14 | 1.8 | Average inhibition $\geq 70\%$ (n=3) |
| Secretion inhibition | 4 | 0.5 | $IC_{50} \geq 50 \mu M$ |
| Translocation inhibition | none | none | $IC_{50} \leq 25 \mu M$ |

Given the small size and composition of the library (776 FDA-approved drugs), it was not surprising that none of these compounds inhibited the assembly of PopD into functional translocons. However, the purpose of utilizing this library was to test the assay under automated high throughput screening conditions and evaluate the strategy for classifying and confirming inhibitors from all the primary hits (Figure 6). This is important because the assay targets the final step in a series of sequential processes (transcription, translation, secretion, etc.) required for effector translocation into the target cell. Overall, our comprehensive evaluation strategy enabled the identification and classification of hits, allowing us to gain insights into their off-target activities and prioritize potential candidates for further investigation.

## Conclusions

The successful implementation of the split NanoLuc technology into the T3SS translocon assembly process, utilizing the PopB-assisted insertion of PopD, allowed the detection of T3SS inhibition. By employing a comprehensive strategy involving secondary and orthogonal assays, it will be possible to find and classify inhibitors that interfere with T3SS secretion, impede T3SS translocation, or impact bacterial growth irrespective of the T3SS. Furthermore, direct monitoring of the exposure of the PopD N-terminus to the host cell cytosol will prove invaluable as a tool for studying the still elusive mechanism of T3SS translocon assembly in *P. aeruginosa* and related pathogens.

## Experimental Section

### Bacterial plasmids and PAK strains construction

The DNA fragment coding for NLuc_10_-GSSGGSSG plus the first 18 amino acids after Met of N-terminus of PopD was synthesized and provided into a DNA shuttle vector pUCIDT (Integrated DNA Technologies). The DNA coding for this fragment was amplified from the shuttle vector using PCR with the following forward (ACGCTTGAGAGGAGACGTCAC) and reverse (GCGATCGGCTCGGAGGGGATC) oligonucleotides. The PCR product was gel purified from a 1.5% Agarose TAE gel using the Zymoclean Gel Extraction Kit (Genesee Scientific). This fragment was then built together with other parts of the pUCP18-GHD vector to obtain the pNLuc10-popD plasmid. This plasmid originated in the pUCP18 vector which contains the *P. aeruginosa* intrinsic pcrG promoter and the coding sequences for both, the PcrH chaperone and PopD [24]. The pUCP18 vector is an *Escherichia-Pseudomonas* shuttle vector [57] and contains the high-copy-number ColE1 origin of replication, the broad-host-range origin of replication oriV which comes from *P. aeruginosa*, and the ampicillin resistant gene (AmpR) for selection. The amino acid sequence for the protein produced by this construct is shown in Figure S1a. The pUCP18-GHNLuc_10_-PopD plasmid was transformed first into the PAKΔD strain to evaluate the functionality of the construct using the cell rounding assay as described previously [24]; and second, into the PAKΔESTYΔD and the PAKΔESTYΔBD strains to be used for the high throughput assay using electroporation as described in [58].

### Stable cell line construction

Construction of the NLuc_1-9_ lentiviral transfer plasmid, and creation of the final NLuc_1-9_-HeLa clonal cell line were completed by the Cell Culture Core Facility at the Institute for Applied Life Sciences, University of Massachusetts-Amherst. The DNA coding for the first 156 amino acids of optimized NanoLuc (NLuc_1-9_) [42] was custom made and provided into the DNA shuttle vector pUCIDT (Integrated DNA Technologies).

The DNA fragment coding for NLuc_1-9_ was amplified from the custom-made shuttle vector using PCR and the following forward (GCTAGCTCGAGGCGGCCGCCATGGTCTTTACCTTGGAGGATTTTGTCG) and reverse (GGTCCAGAACTGATATTATCTATGCGG) oligonucleotides. The PCR product was digested with BamHI (NEB), and the resulting 514 bp fragment was gel purified from a 1.5% Agarose gel using the GeneJet Gel Extraction Kit (Thermo Scientific). The plasmid pLenti-DsRed_IRES_EGFP (Addgene plasmid # 92194) was cleaved with AfeI and BamHI endonucleases (NEB) to remove the DsRed gene. The larger fragment of the vector was gel purified as described above and the two fragments were ligated by T4 ligase (NEB) to generate the pLenti-NLuc_1-9__IRES EGFP transfer plasmid. Lenti-X 293T cells (TaKaRa), 8 x 10^5^ cells per well of a 6 well dish, were plated a day prior to transfection in 3 mL DMEM (Gibco), 10% FBS (Corning). Overnight (∼16 h) cultures were transfected with lentiviral packaging plasmids pMD2.G and psPAX2 (Addgene plasmid # 12259 and 12260, respectively) and transfer plasmid pLenti-NLuc_1-9__IRES_EGFP (Figure S5) using Lipofectamine2000 (Invitrogen). DNA ratio of plasmid mix was 2:1:4, respectively, with a combined total of 3 µg per well. Lipofectamine2000 per well totaled 9 µL. Transfections were incubated overnight. Transfection mixes were removed, and 3 mL fresh growth media was added per well. Cultures were incubated for additional 48 h. Cell culture supernatants containing lentivirus were collected, passed through a 0.45 µm filter, aliquoted and stored −80°C. HeLa cells (ATCC CCL-2) were added to wells of a 48 well plate; 4 x 10^4^ in 0.4 mL MEM (Lonza), 10% FBS (Corning), and incubated overnight. Media was replaced with 0.2 mL of NLuc_1-9_-IRES GFP lentivirus dilution (1:5, 1:25, 1:125, or 1:625) and incubated overnight. 0.2 mL of growth media was added per well, then incubated for an additional 48 h. On day 3, following transduction, GFP expression was evaluated by flow cytometry. Single cell clones were further isolated. The sequence for the NLuc_1-9_ produced in this cell line is shown in Figure S1b.

### PopD assembly assay

NLuc_1-9_-HeLa cells were maintained in a T25 flask (Corning) with DMEM (Hyclone) plus 10% FBS (Hyclone), incubated at 37°C, 5% CO_2_ in the incubator and when needed, the cells were seeded in white 96 well plate (flat bottom, tissue culture treated), each well containing ∼10,000 cells. Corresponding PAK strains were growing overnight with corresponding antibiotics in Lysogeny broth (LB), Miller’s (Fisher) at 37°C, 180 rpm. Cultures were then diluted into OD_600_ of 0.1 in new fresh LB medium and incubated until OD_600_ reached 0.5-1.0. NLuc_1-9_-HeLa cells were then washed twice with DPBS with calcium and magnesium, and 100 µl DMEM without FBS was added in each well. Then the NLuc_1-9_-HeLa cells were infected with corresponding PAK strains at MOI = 50, and incubated at 37°C, 5 % CO_2_ for 45 min. NLuc_1-9_- HeLa cells were washed twice with DPBS to remove bacteria and DMEM medium, and finally 75 µl Opti-MEM (Gibco) were added into each well. DMEM is replaced with Opti-MEM to eliminate the phenol red that interferes with the luminescence signal (Figure S4). Cells were incubated at 37°C, 5 % CO_2_ for an additional 30 min to allow NanoLuc complementation. Next, 25 µl Nano-Glo live cell reagent (Promega) was added to each well, and the plate was placed on shaker (80 rpm) at room temperature (∼20-25 °C) for 10 min. Luminescence signal was collected at Synergy H1 plate reader. Measurements were done using default luminescence fiber mode, no filter, read height of 7 mm, and a gain set at 135. The signal was recorded for 15 min at intervals of 1 min, and the average of the last 5 points was used as the luminescence signal of each sample.

### Spectroscopic measurements

Fluorescence measurements were made with a Fluorolog-3 photon-counting spectrofluorimeter as reported earlier [23]. For mCherry emission maxima λ_max_ measurements, the wavelength for mCherry excitation was set to 585 nm and emission was scanned from 595 nm to 700 nm, every 1 nm using 1s integration time. The bandpass was 2 nm for excitation and 4 nm for emission. For NanoLuc reaction light emission was scanned from 400 nm to 500 nm, every 1 nm using 1s integration time. The bandpass was 4 nm for emission. Absorbance measurements were done using a Beckman DU800 spectrophotometer. The absorbance was scanned from 400 nm to 500 nm, every 1 nm using 0.1s integration time.

### Z’-factor, and S/N determination

For the manual assay, the luminescent signal of nine replicas for inhibition negative control (functional translocon) and inhibition positive control (non-functional translocon using a ΔpopB-based strain) were measured as described above for the split NanoLuc assay. The Z’-factor was calculated using equation 1.

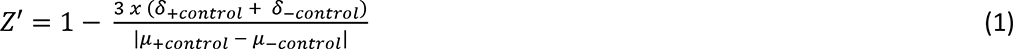

where δ is the standard deviation and µ is the mean of the measured replicas. For automated assays the Z’ was calculated using a total of 144 replicas in duplicate for each positive (plus MBX 2401) and negative (plus DMSO) control.

The S/N ratios were calculated using equation 2. The S/N ratio is only an indication of the degree of confidence with which a signal can be regarded as real, i.e., different from the associated background noise. In this assay the signal to be distinguished from the background noise is the positive inhibition control or lack of signal. The background noise is provided by the luminescence that results from the assembly of variable number of functional translocons on cells expressing variable amounts of NLuc_1-9_.

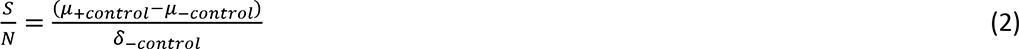

The S/N ratio should not be confused with the signal to background (S/B) ratio. The S/B ratio calculated using equation 3 does not contain any information regarding data variation.

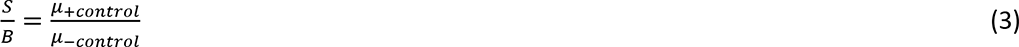

Equations 1, 2, and 3 were used as defined by Zhang et al. [47].

### PopD assembly assay on 96 well plate

NLuc_1-9_-HeLa cells were grown in T75 culture flasks (Fisher) in DMEM with 10% FBS at 37°C, 5% CO_2_ in the incubator. In 96-well white, flat-bottomed TC treated microplates, 10,000 cells were seeded per well in 100 µL of DMEM with 10% FBS using a MultiDrop Combi reagent dispenser (Thermo Scientific). Plates were incubated overnight at 37°C, 5% CO_2_.

PAKΔESTYΔD: NLuc_10_-PopD was grown in 10 mL of LB Miller supplemented with 100 µg/mL carbenicillin. Cultures were incubated at 37°C, 200 rpm overnight. The overnight culture was diluted to an OD_600_ = 0.1 in 10 mL LB Miller supplemented with 100 µg/mL carbenicillin. The bacterial culture was incubated at 37°C, 200 rpm until it reached an OD_600_ = 0.6-0.8 (∼1.5 h).

NLuc_1-9_-HeLa cells were washed twice with 100 µL DPBS (with calcium and magnesium; pre-warmed) using a ELx405 Select CW microplate washer (BioTek Instruments). With the Combi speed set to low, 100 µL of fresh DMEM without FBS was added to each well. Using a Caliper LifeSciences SciClone ALH 3000 liquid handler equipped with a 96-tip pipetting head, 1 µL of compound was transferred from compound screening plates to assay plates, giving a final DMSO concentration of 1% (v/v) and final compound concentration of 25 µM. Only DMSO was added to the first column of the assay plate (negative control), whereas the T3SS inhibitor MBX-2401 (10 µM final concentration) was added to the last column of the assay plate (positive control).

The NLuc_1-9_-HeLa cells were then infected with the PAK strain at MOI = 50. The MOI was calculated using (CFU/ml) = 2.0087 × (OD_600_) + 0.0764 [59]. The bacterial culture is then diluted in DMEM (no FBS) such that 10 µL of bacteria is added per well. Bacteria were added using a multichannel pipette. Plates were centrifuged at 1200 rpm for 1 minute to facilitate the interaction of the bacteria and the HeLa cells. Infection took place for 45 minutes at 37°C in 5% CO_2_. After infection, NLuc_1-9_-HeLa cells were washed twice with pre-warmed 100 µL DPBS (with calcium and magnesium) using the microplate washer to remove bacteria. With the Combi speed set to low to minimize cell detachment, 90 µL of Opti-MEM was added to each well and plates were incubated for 30 minutes at 37°C in 5% CO_2_ to allow complete NanoLuc complementation. Next, 10 µL Nano-Glo live cell reagent (Promega) were added to each well using the Combi. Plates were covered with aluminum seals and placed on a shaker at room temperature for 20 minutes to stabilize the signal. Luminescence was then measured on the Envision plate reader.

% inhibition was calculated using equation 2

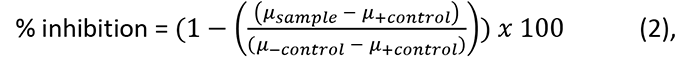

where µ is the mean of the measure replicas.

### High throughput screening testing

To evaluate the assays’ readiness for high throughput, a pilot screen against an FDA-approved compound library, n = 776 compounds was conducted. The assays were performed in 96 well plate format using automation equipment, following the protocol described above, with eleven plates tested in total. DMSO was used as the negative control and MBX-2401, a T3SS inhibitor, was used as a positive control. The control values were like those observed during the assay development stage, and the controls on each assay plate demonstrated good plate-to-plate reproducibility as described in Results. The final concentration of FDA compounds assessed in the assay was 25 μM.

### *P. aeruginosa* T3SS secretion assay

T3SS gene regulation is tightly coupled to ExsE secretion. A *P. aeruginosa* strain was previously constructed with the luxCDABE operon of *Photorhabdus luminescens* transcriptionally fused to the Type 3 effector gene *exoT* (MDM-852). Under T3SS-inducing conditions, ExsE is secreted, and luminescence is observed, while when secretion is inhibited, luminescence is significantly reduced. Inhibition of T3SS-mediated secretion by *P. aeruginosa* MDM-852 was measured in black 96-well microplates as previously described [54]. In brief, a 5 mL culture of *P. aeruginosa* MDM-852 was grown in LB Miller supplemented with 10 µg/mL gentamicin (LB Miller Gm10) and incubated overnight at 37˚C with aeration. An aliquot of 100 µL from the overnight culture was diluted into 10 mL of fresh LB Miller Gm10 and incubated at 37˚C with aeration for approximately 2.5 h.

A dilution plate with the compounds of interest was prepared, with the highest concentration of compound at 50-fold the desired starting concentration for the assay. For the single point secretion inhibition assay, the compound plate was prepared such that compounds would be evaluated at a final concentration of 25 µM, whereas for the dose response assay, serial half dilutions were performed from the starting concentration of 50 µM. 2 µL of compound was added from the dilution plate in triplicate to the assay plate (two compounds were tested in triplicate per assay plate).

After 2.5 h the OD_600_ was read, and the subculture was diluted to a final OD_600_ of 0.05 in LB Miller Gm10. The culture is aliquoted into two falcon tubes, one to be treated with ethylene glycol-bis(β-aminoethyl ether)-N,N,N′,N′-tetraacetic acid (EGTA) and one without. EGTA (0.5 M, pH 8.0) was added to a final concentration of 3 mM. One hundred µL was dispensed into columns 1-11. One hundred µL of the remaining back-diluted culture (no EGTA) was added to column 12 of the assay plate.

The plates were incubated for 5 h at room temperature in the dark. After incubation, luminescence was measured using an Envision Multilabel microplate reader (PerkinElmer). The IC_50_ value from each compound replicate on the assay plate was calculated in GraphPad Prism 9.0.

### Inhibition of *P. aeruginosa* growth assay

MIC determination was done by the broth microdilution method described in the CLSI (formerly NCCLS) guidelines [53] and expressed in µM to facilitate comparisons with secretion IC_50_ values. For the single point growth inhibition assay, the compound plate was prepared such that compounds would be evaluated at a final concentration of 25 µM, whereas for the dose response MIC assay, serial half dilutions were performed from the starting concentration of 50 µM.

### T3SS effector translocation assay

CHO cells were incubated with *P. aeruginosa* and each compound as described [55]. Cell death was measured by lactate dehydrogenase (LDH) release to detect T3SS effector ExoU-mediated lysis of CHO cells. Percent cell death (expressed as percent LDH release) was calculated relative to that of the uninfected control (0%), and that of cells infected with *P. aeruginosa* unprotected by test compound (100%). The same assay was run in parallel with inhibitor and CHO cells, but without *P. aeruginosa* cells, to evaluate the CHO cell cytotoxicity (CC_50_) of each compound.

### T3SS functionality based on cell rounding assay

The cell rounding assay was performed as reported earlier [41]. Briefly, corresponding PAK strains were incubated for 120 min with HeLa cells and the T3SS functionality was evaluated by the rounding of the HeLa cells observed at different intervals caused by translocated effectors on actin cytoskeleton.

### Supporting Information

Figure S1 Primary sequence of proteins and peptide used in the assay; Figure S2 Variability of the screening of the FDA compound library Replica 2; Figure S3 Determining the cutoff for the screening assay; Figure S4 Spectra of the furimazine luminescence and potential interferences; Figure S5 Scheme of the plasmid used in stable cell line construction.

### Abbreviations Used

T3SS, Type III Secretion System; PAK, *P. aeruginosa* PAK strain; MOI, Multiplicity Of Infection; NLuc, NanoLuc luciferase; MIC, Minimal Inhibitory Concentration; LB, Lysogeny Broth; RLUs, Relative Light Units; CBD, Cannot Be Determined.

### Author contributions

^#^ H.G. and E.G contributed equally to this work. H.G. constructed the plasmids, bacterial and cell strains, and set up the experimental conditions for the assay. E.G. adapted the assay to high throughput automatic measurements, and performed the screen using the compound library. H.G. and E.G. performed secondary assays to confirm hits. A.P.H. and T.J.O. designed the project and supervised research. All authors analyzed the data and prepared the manuscript.

## Acknowledgements

This work was supported in part by UMass IALS Midigrant and the UMass Faculty Research Grant/Healey Endowment Grant (to A.P.H.), and by a National Institutes of Health Grant R41 AI149922 (to T.J.O. and A.P.H.). Stable NLuc_1-9_-HeLa cell clone was obtained by the University of Massachusetts Cell Culture Core Facility (RRID: SCR_023477).

