## Supplemental Information for "Cell-based assay to determine Type 3 Secretion System translocon assembly in *Pseudomonas aeruginosa* using split luciferase"

**SUPPORTING INFORMATION**

**a.** **NLuc10-PopD**

```
         10        20        30        40        50        60        70        80        90       100
....|....|....|....|....|....|....|....|....|....|....|....|....|....|....|....|....|....|....|....|
MVSGWRLFKKISGSSGGSSGIDTQYSLAATQAAIPSEPIAPGAAGRSVGTPQAAADLPQVPAARADRVELNAPRQVLDPVRMEAAGSELDSSVELLLILF

        110       120       130       140       150       160       170       180       190       200
....|....|....|....|....|....|....|....|....|....|....|....|....|....|....|....|....|....|....|....|
RIAQKARELGVLQRDNENQSIIHAQKAQVDEMRSGATLMIAMAVIAGVGALASAVVGSLGALKNGKAISQEKTLQKNIDGRNELIDAKMQALGKTSDEDR

        210       220       230       240       250       260       270       280       290       300
....|....|....|....|....|....|....|....|....|....|....|....|....|....|....|....|....|....|....|....|
KIVGKVWAADQVQDSVALRAAGRAFESRNGALQVANTVIQSFVQMANASVQVRQGESQASAREGEVNATIGQSQKQKVEDQMSFDAGFMKDVLQLIQQYT

        310
....|....|....
QSHNQAWRAAAGVV
```

**b.** **NLuc1-9**

```
         10        20        30        40        50        60        70        80        90       100
....|....|....|....|....|....|....|....|....|....|....|....|....|....|....|....|....|....|....|....|
MVFTLEDFVGDWEQTAAYNLDQVLEQGGVSSLLQNLAVSVTPIQRIVRSGENALKIDIHVIIPYEGLSADQMAQIEEVFKVVYPVDDHHFKVILPYGTLV

        110       120       130       140       150       160
....|....|....|....|....|....|....|....|....|....|....|....|....|....
IDGVTPNMLNYFGRPYEGIAVFDGKKITVTGTLWNGNKIIDERLITPDGSMLFRVTINSGGRSR
```

**Figure S1 Primary sequence of proteins and peptide used in the assay. a)** Primary sequence of $NLuc_{10}$-PopD. The optimized $NLuc_{10}$ is highlighted in black and GSSGGSSG linker is highlighted in orange. **b)** Primary sequence of optimized $NLuc_{1-9}$ used in the assay.

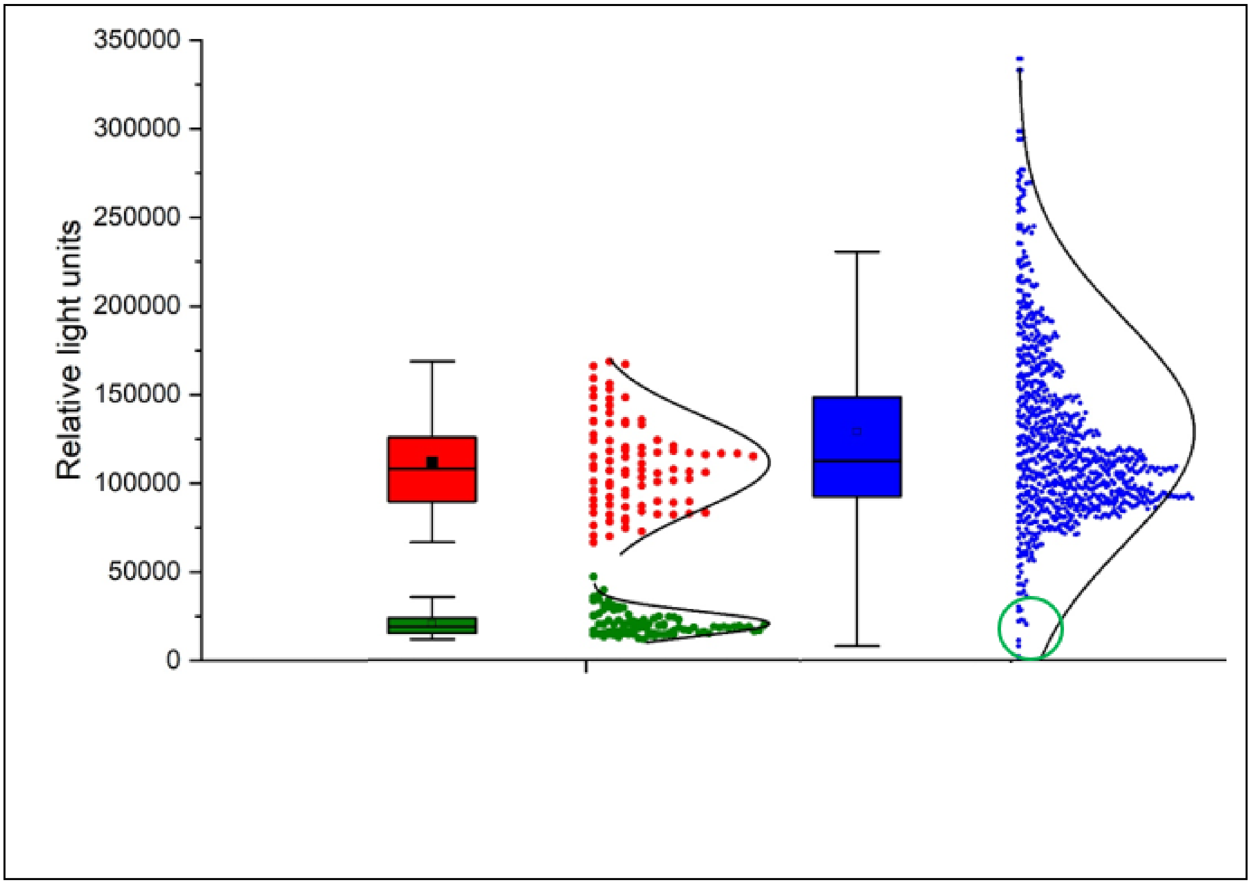


**Figure S2 Variability of the screening of the FDA compound library Replica 2.** Raw data in relative light units were plotted as a box plot with normal distribution for replica 2. The negative control (DMSO-treated) is shown in red, the positive control (MBX-2401) is shown in green, and screened compounds are shown in blue. The average S/B was 5.4, with a Z' value of -0.11 across eleven 96 well plates. The box plot shows the mean (black square), the median (line), the 25-75 percentage range (boxes), and the standard deviation (whisker). The green circle shows the compounds selected as hits.

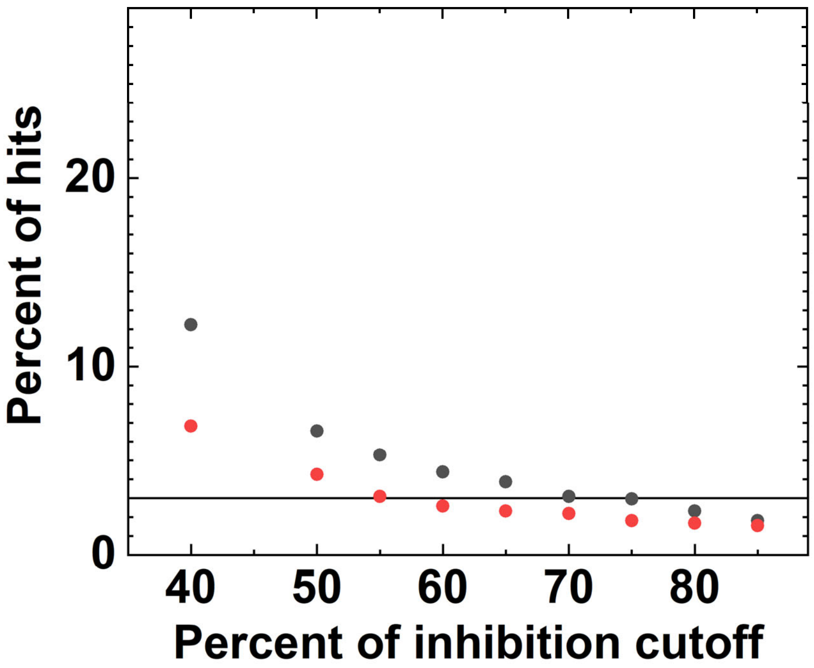


**Figure S3**. **Determining the cutoff for the screening assay.** The percentage of hits was plotted as a function of the percentage of inhibition cutoff set for the replicas. Three percent or less hits (black line) were obtained when the cutoff was set above 70% inhibition.

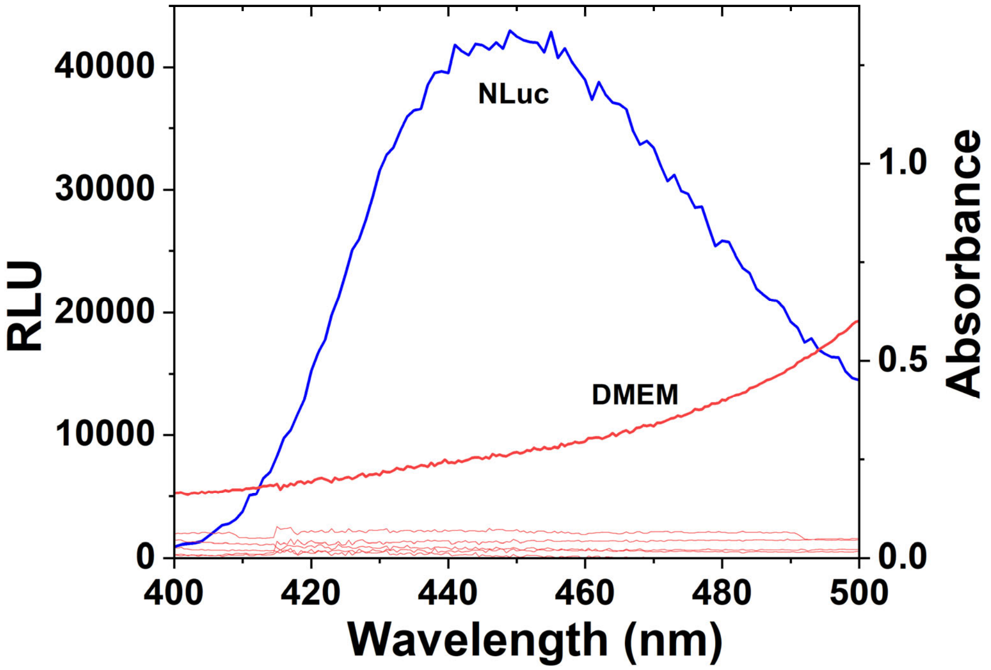


**Figure S4 Spectra of the furimazine luminescence and potential interferences.** Emission spectrum for the light emitted by NanoLuc catalysis of furimazine is shown in blue (left axis). Absorbance spectra for DMEM, Opti-MEM, 25% furimazine, MBX4918 100 µM, MBX2401 100 µM, and 5% DMSO are shown in red (right axis). Only DMEM medium (indicated) showed significant absorbance at these wavelengths.

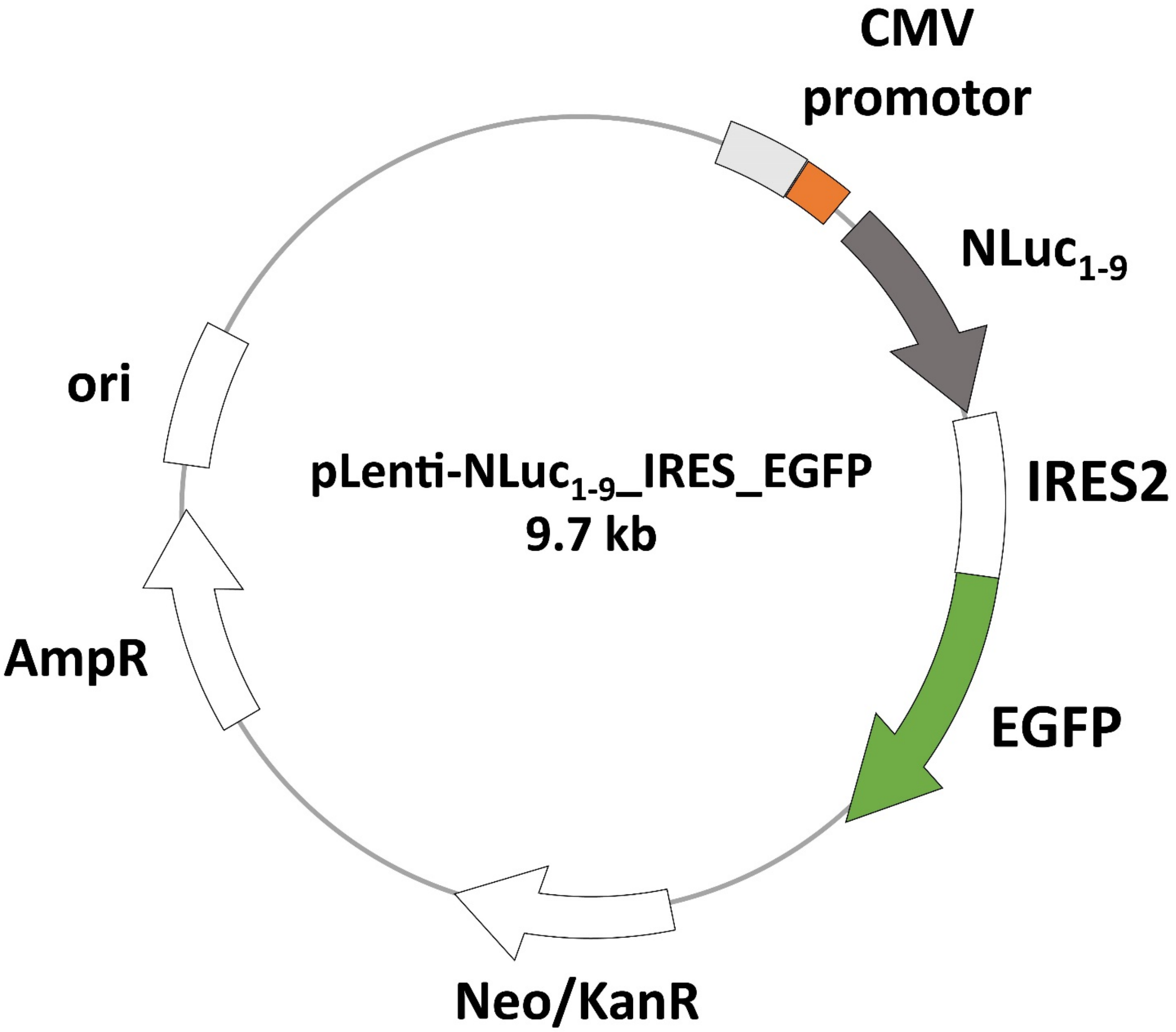


**Figure S5 Scheme of the plasmid used in stable cell line construction.** The CMV promoter (enhancer in light gray) regulates the expression of the $NLuc_{1\text{-}9}$ (highlighted in dark gray) and the eGFP (highlighted in green) is co-expressed under the same CMV promoter with the help of the Internal Ribosome Entry Site 2 (IRES2) sequence. The plasmid contains origin of replication (ori) from *E.coli* and Ampicillin resistant gene (AmpR) for bacterial selection. The plasmid also contains Neomycin/Kanamycin resistant gene (Neo/Kan R) for later selection in mammalian cells.
